# Accurate detection and identification of insects from camera trap images with deep learning

**DOI:** 10.1101/2022.10.25.513484

**Authors:** Kim Bjerge, Jamie Alison, Mads Dyrmann, Carsten Eie Frigaard, Hjalte M. R. Mann, Toke Thomas Høye

## Abstract

Reported insect declines have dramatically increased the global demand for standardized insect monitoring data. Image-based monitoring can generate such data cost-efficiently and non-invasively. However, extracting ecological data from images is more challenging for insects than for vertebrates because of their small size and great diversity. Deep learning facilitates fast and accurate insect detection and identification, but the lack of training data for coveted deep learning models is a major obstacle for their application. We present a large annotated image dataset of functionally important insect taxa. The primary dataset consists of 29,960 annotated insects representing nine taxa including bees, hoverflies, butterflies and beetles across more than two million images recorded with ten time-lapse cameras mounted over flowers during the summer of 2019. The insect image dataset was extracted using an iterative approach: First, a preliminary detection model identified candidate insects. Second, candidate insects were manually screened by users of an online citizen science platform. Finally, all annotations were quality checked by experts. We used the dataset to train and compare the performance of selected You Only Look Once (YOLO) deep learning algorithms. We show that these models detect and classify small insects in complex scenes with unprecedented accuracy. The best performing YOLOv5 model consistently identifies nine dominant insect species that play important roles in pollination and pest control across Europe. The model reached an average precision of 92.7% and recall of 93.8 % in detection and classification across species. Importantly, when presented with uncommon or unclear insects not seen during training, our model detects 80% of individuals and usually interprets them as closely related species. This is a useful property to (1) detect rare insects for which training data are absent, and (2) generate new training data to correctly identify those insects in future. Our camera system, dataset and deep learning framework show promising results in non-destructive monitoring of insects. Furthermore, resulting data are useful to quantify phenology, abundance, and foraging behaviour of flower-visiting insects. Above all, this dataset represents a critical first benchmark for future development and evaluation of deep learning models for insect detection and identification.

## 1. Introduction

Multiple lines of evidence suggest widespread declines in the abundance, diversity, and biomass of insects (1). Standardised insect monitoring is needed to unravel the magnitude of these trends across taxa, regions, and time periods (2). In particular, automated sensor systems (e.g. insect camera traps) and data extraction algorithms (e.g. computer vision and deep learning) are valuable tools to understand trends in insects and their underlying drivers (3–6). However, the potential of deep learning to process such sensor signals (e.g. detecting insects in images) for insect monitoring hinges upon (1) choice and positioning of sensors; (2) accuracy, speed and usability of deep learning models and (3) quantity, quality and accessibility of training data.

Within deep learning and computer vision, convolutional neural networks (CNNs) are readily applied in ecological research (7) and are particularly useful for detection and classification tasks (8). Studies have demonstrated considerable advances in use of CNNs for insect identification and monitoring; modern cameras offer sufficient resolution to identify many insect species, while CNNs can automate species classification with very high accuracy (9). However, many barriers remain, and the availability of standardised training data is a major bottleneck to wider use and application (4). Indeed, most rigorous tests of CNNs in entomology have been based on dead insects (10), where curated collections of specimens of known identity are most readily available (11), or non-standardised images from citizen science databases (12).

A particularly exciting prospect enabled by deep learning is automated, non-invasive monitoring of insects and other small organisms in their natural environment (13). However, this brings specific challenges that reach beyond the classification tasks addressed in previous studies (11,12). Most importantly, small insects must be detected and localised within complex and dynamic scenes of natural vegetation. While detection and classification of insects *in situ* can be achieved, the above factors can lead models to falsely identify background elements as objects of interest (14). Furthermore, insects often exhibit high intra-class variation (e.g. a butterfly species with wings open vs. closed) and several individuals from multiple, similar species groups may appear simultaneously within an image (15). Models trained on museum specimens or opportunistic image libraries are not equipped to localise and identify small insects on natural vegetation. Training images must include relevant insect taxa, but also relevant backgrounds against which to detect them; otherwise, models will generate large numbers of false positive detections, resulting in low model precision.

Another challenge in automated insect monitoring is the inevitable appearance of species or classes of insects for which adequate training data do not exist. The immense diversity of insects, as well as the unevenness of ecological communities, mean that it is not feasible to obtain training data to perfectly classify all species. Thus, as is the case with DNA reference libraries, scientists will need to work with incomplete and imbalanced image reference libraries for the foreseeable future. In practice, models will need to be retrained when images of poorly represented species or classes become available. In the meantime, most CNNs can either (1) fail to detect the unfamiliar insect (i.e. classify it as background) or (2) detect the unfamiliar insect and falsely attribute it to a similar, trained class. Both of these outcomes are imperfect, resulting from practical limitations in both training data and model specification. However, detecting unfamiliar insects and interpreting them as close relatives is advantageous if model predictions are regularly screened; in this way novel classes can be identified, labelled, and incorporated into future training datasets. Furthermore, many important ecological questions are not sensitive to minor misidentifications, concerning e.g. function, activity, abundance or biomass of broad taxonomic groups. Clearly, the way that insect detection and classification models respond to novel and potentially nested classes demands further investigation (16).

Insects move quickly and their interactions with the environment, such as pollination events, are ephemeral. Thus, for image-based monitoring, a high image capture frequency is preferable. Image processing with a deep learning model can either be applied in real-time (i.e. in parallel with image capture) or images can be stored and processed subsequent of collection. For detecting objects in real-time or processing large amount of data, computational speed is critical, and the open source deep learning network, You Only Look Once (YOLO), which is known to be particularly fast has gained strong support (17). YOLO (17) is a unified object detection architecture for deep neural networks where YOLOv5 is written in Python and use PyTorch as the machine learning framework. YOLOv5 can process images and detect objects with a frame rate ranging from 30 to 100 frames per second depending on the model size, measured by the number of trainable parameters, the size of the image, and the computers graphical processing unit (GPU).

Here, we present the data and research infrastructure for automated image-based insect monitoring and make it freely available. Specifically, we provide a remarkable training dataset and validated deep learning models to demonstrate accurate, automated detection and classification of insects in complex ecological scenes. We developed and used a custom-built time-lapse camera system to capture images of plants and insects. From the images, we created the datasets for training insect detection models (YOLO) which could be deployed for on-board processing and thus real-time presentation of insect monitoring data. We demonstrate the optimal configuration to detect and classify nine dominant insect taxa, and show abundance of those taxa through an entire summer comprising more than two million images. Furthermore, we evaluate how models perform when presented with insects not confirmed to fall within those nine species, simulating the scenario where unclear specimens or rare species are encountered during monitoring. Finally, we demonstrate how the trained model can be used to monitor the seasonal abundance of insect species.

## 2. Results

During the period June to September 2019, ten cameras (Figure *1*) were in operation on a green roof with a mix of 13 species of *Sedum* plants (18). The *Sedum* plant species flower at different times during the growing season, so there was a continuous availability of flowers during the recording period from 24. June to 30. September. Data from the first six weeks were unfortunately lost from two cameras due to rainwater entering the hard drive in one system and destroying it. In total 2,347,944 images from the 10 cameras were collected during the study, occupying 591 GB of data. Examples of collected images are shown in Figure 2.

**Figure 1:**
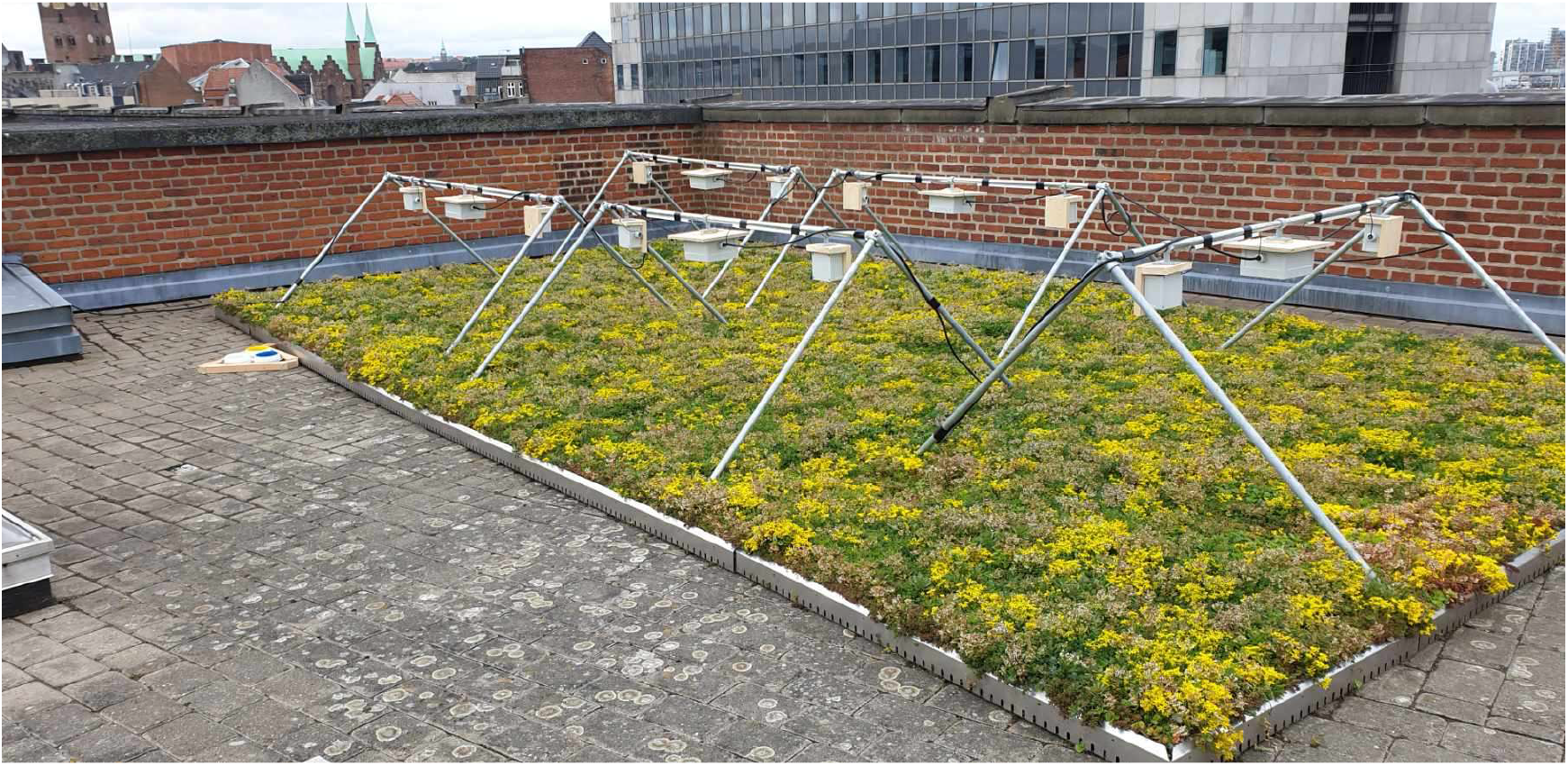
Five insect monitoring systems with ten cameras mounted on a roof with Sedum plants. The system collects time-lapse images with an interval of 30 seconds during the daytime. The white central box on each frame contains a Raspberry Pi computer with an external hard disk connected to two Logitech web cameras. The system is powered through the power grid.

**Figure 2:**
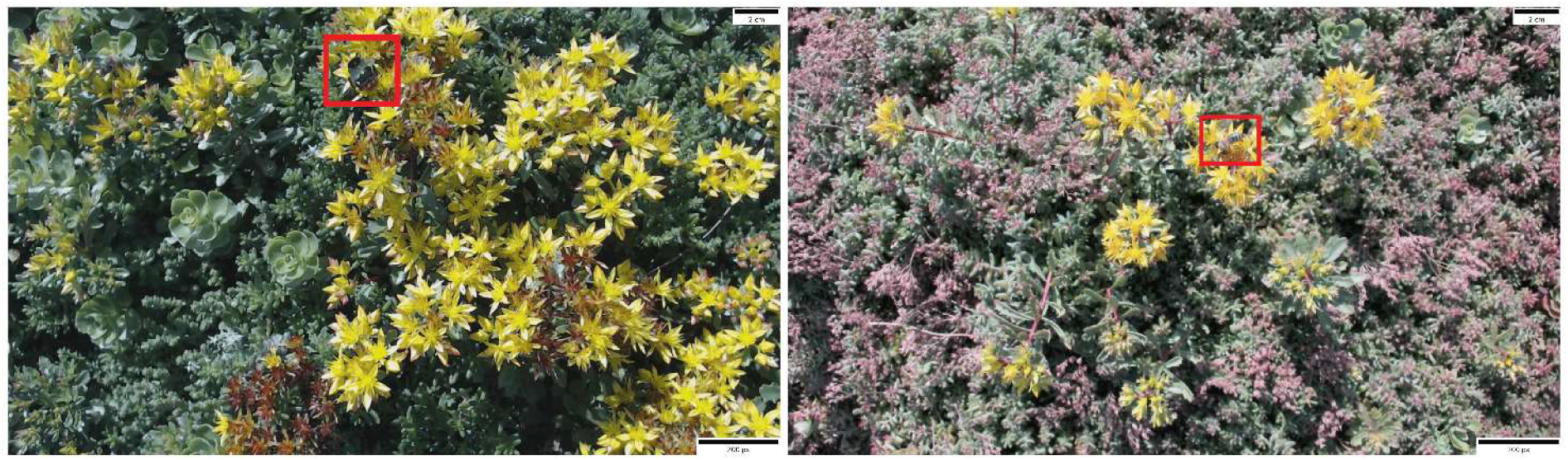
Images of *Sedum* plants with one bumblebee (left) and honey bee (right) highlighted by the red rectangle. The image size is 1920×1080 pixels and covers a camera view of approximately 35 × 22 cm.

An overview of the initial and final training and validation datasets, as well as the additional test dataset, are shown in Table 1. Example images of individual insect species for each class are shown in Figure 3. Different classes are not evenly represented in the data, with the smallest sample sizes for species such as *Aglais urticae* and *Eristalis tenax*. Background images comprise a selection of images from the ten cameras representing the growing season of *Sedum* plants, with different flowering colours, but without insects. Table 1 shows an overview of the occurrence of all species in the annotated dataset for training, validation and testing of the CNN algorithm.

**Table 1:**
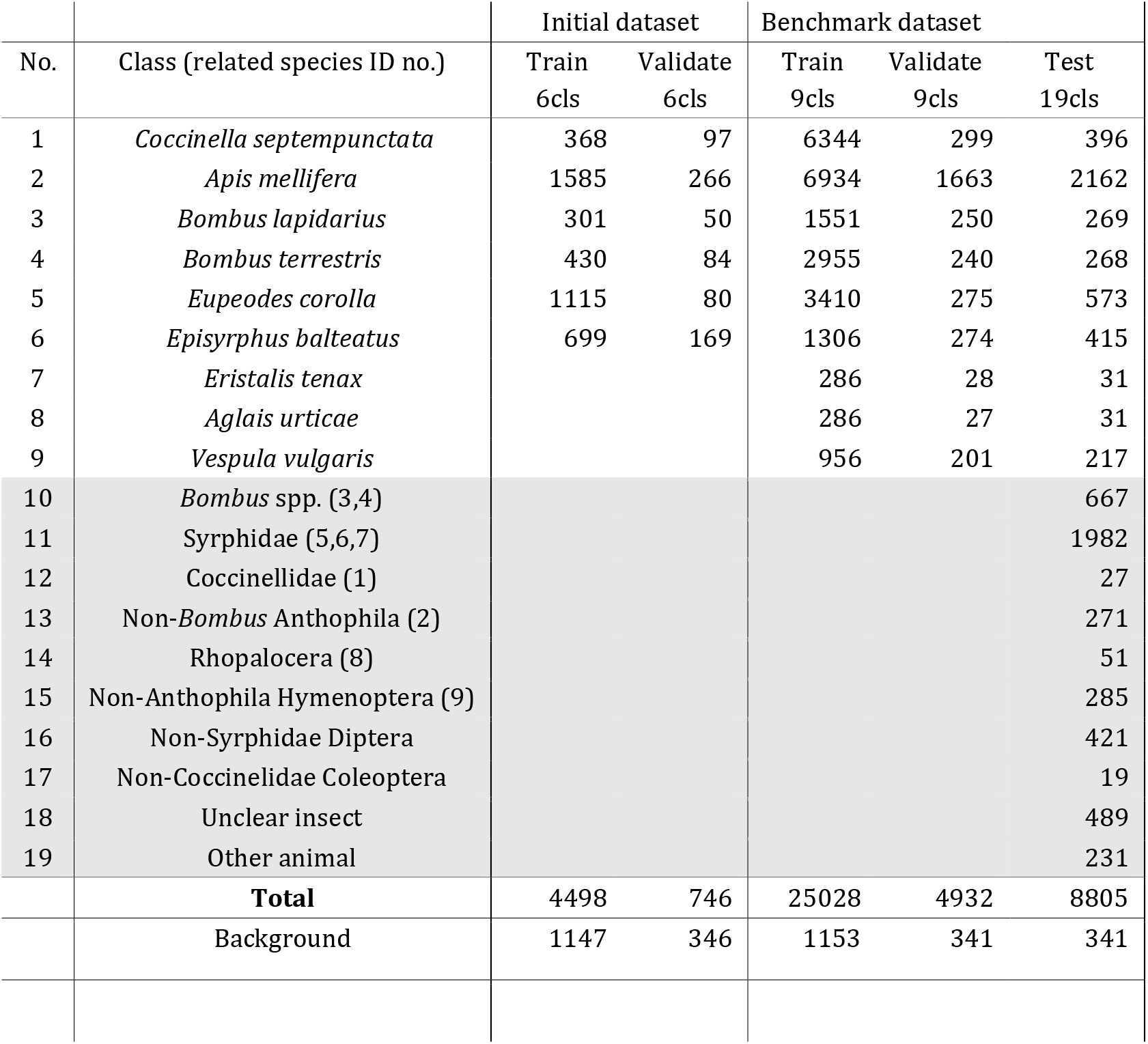
Number of insect classes in the dataset used for training, validation and test. Insects were annotated from the images with *Sedum* plants. The background class contains images without any insects. A train and validation dataset with six species (6cls) was used to train the first YOLO model. A benchmark training and validation dataset with an approximate split percentages of 85/15 with nine species (9cls) was used to train the final evaluated YOLO models. The test component of the benchmark dataset contains additional unfamiliar classes 10-19 that contain other species of bees, flies and beetles, unclear insects and other animals. Classes 1-9 are listed in brackets next to their closest relative of the unfamiliar classes 10-19.

| No. | Class (related species ID no.) | Initial dataset |  | Benchmark dataset |  |  |
| --- | --- | --- | --- | --- | --- | --- |
|  |  | Train<br>6cls | Validate<br>6cls | Train<br>9cls | Validate<br>9cls | Test<br>19cls |
| 1 | *Coccinella septempunctata* | 368 | 97 | 6344 | 299 | 396 |
| 2 | *Apis mellifera* | 1585 | 266 | 6934 | 1663 | 2162 |
| 3 | *Bombus lapidarius* | 301 | 50 | 1551 | 250 | 269 |
| 4 | *Bombus terrestris* | 430 | 84 | 2955 | 240 | 268 |
| 5 | *Eupeodes corolla* | 1115 | 80 | 3410 | 275 | 573 |
| 6 | *Episyrphus balteatus* | 699 | 169 | 1306 | 274 | 415 |
| 7 | *Eristalis tenax* |  |  | 286 | 28 | 31 |
| 8 | *Aglais urticae* |  |  | 286 | 27 | 31 |
| 9 | *Vespula vulgaris* |  |  | 956 | 201 | 217 |
| 10 | *Bombus* spp. (3,4) |  |  |  |  | 667 |
| 11 | Syrphidae (5,6,7) |  |  |  |  | 1982 |
| 12 | Coccinellidae (1) |  |  |  |  | 27 |
| 13 | Non-*Bombus* Anthophila (2) |  |  |  |  | 271 |
| 14 | Rhopalocera (8) |  |  |  |  | 51 |
| 15 | Non-Anthophila Hymenoptera (9) |  |  |  |  | 285 |
| 16 | Non-Syrphidae Diptera |  |  |  |  | 421 |
| 17 | Non-Coccinellidae Coleoptera |  |  |  |  | 19 |
| 18 | Unclear insect |  |  |  |  | 489 |
| 19 | Other animal |  |  |  |  | 231 |
|  | **Total** | 4498 | 746 | 25028 | 4932 | 8805 |
|  | Background | 1147 | 346 | 1153 | 341 | 341 |

**Figure 3:**
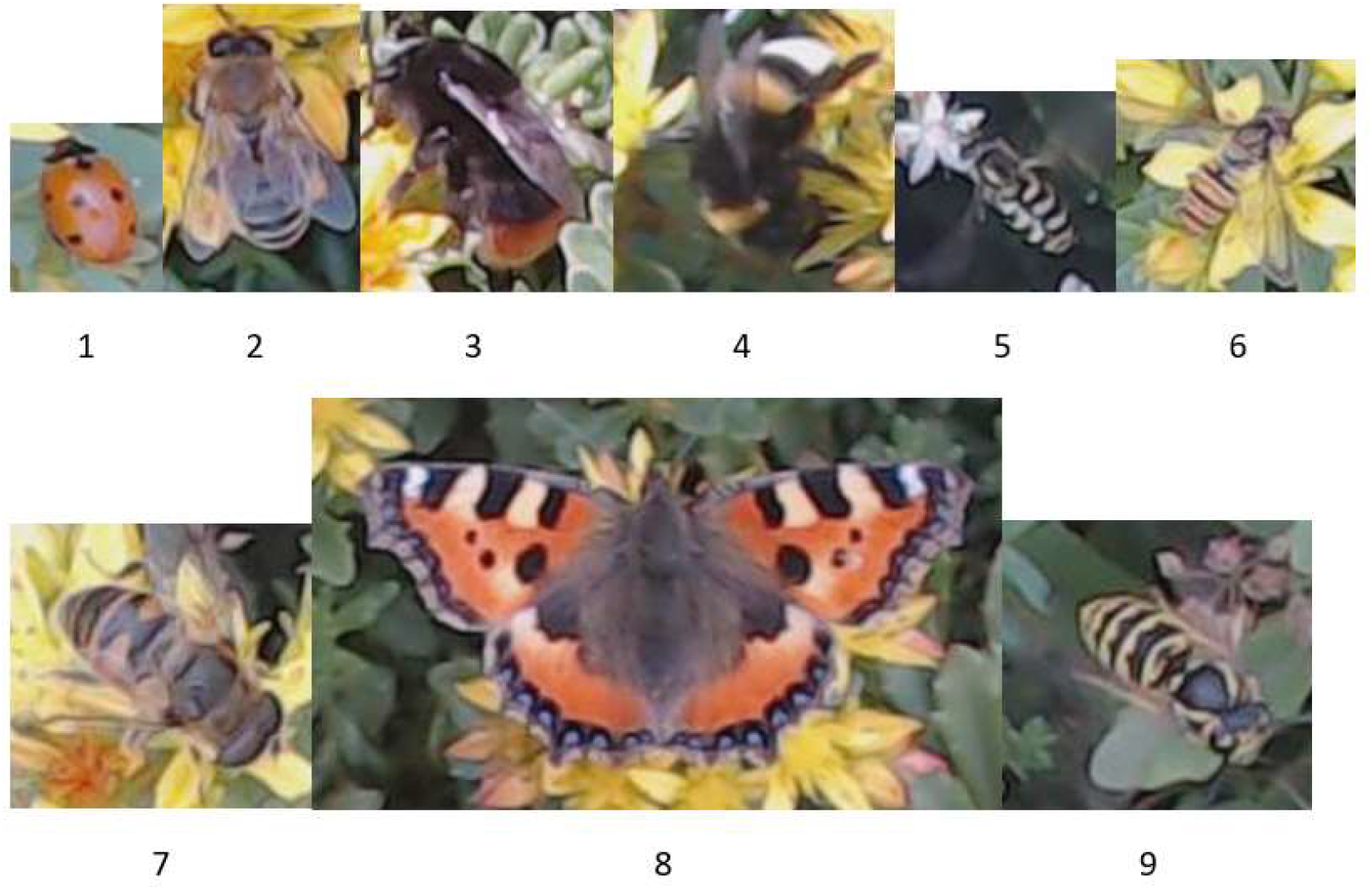
Examples of the common insect species that were observed and labelled for training and validation of the YOLO models. The numbers refer to the species listed in Table 1.

We trained a total of 10 different YOLO models as listed in Table *2* based on the datasets listed in Table *1*. No. 1 is the initial YOLOv3 model trained on the small dataset with six classes, which we used to create the benchmark dataset with nine classes. No. 2 and 3 are two different YOLOv5 models trained on the initial (6 classes) dataset with varying image and model size. No. 4 is a YOLOv3 model trained on the large dataset with nine classes to assess the performance gain of adding data for training YOLOv3 models. Model No. 4 also allowed us to compare a YOLOv3 model with different YOLOv5 models all trained and validated on the benchmark dataset. The different YOLOv5 models varied in image and model size (model No. 5 to No. 10).

In Figure *4* the class, object and box losses are shown for training YOLOv5 models with nine classes. For an image size of 640 (640v5m), the validation class loss starts to increase in the area of 275 epochs. For an image size of 1280, object loss starts to increase in the area of 350 epochs. This indicates the start of overfitting and it is before or in this area that the best models should be found when comparing with Figure 5. In Figure 5 the mAP@.5:.95 and F1-scores are shown as a function of epochs for validating the trained models and red marks indicates the best models found.

**Figure 4:**
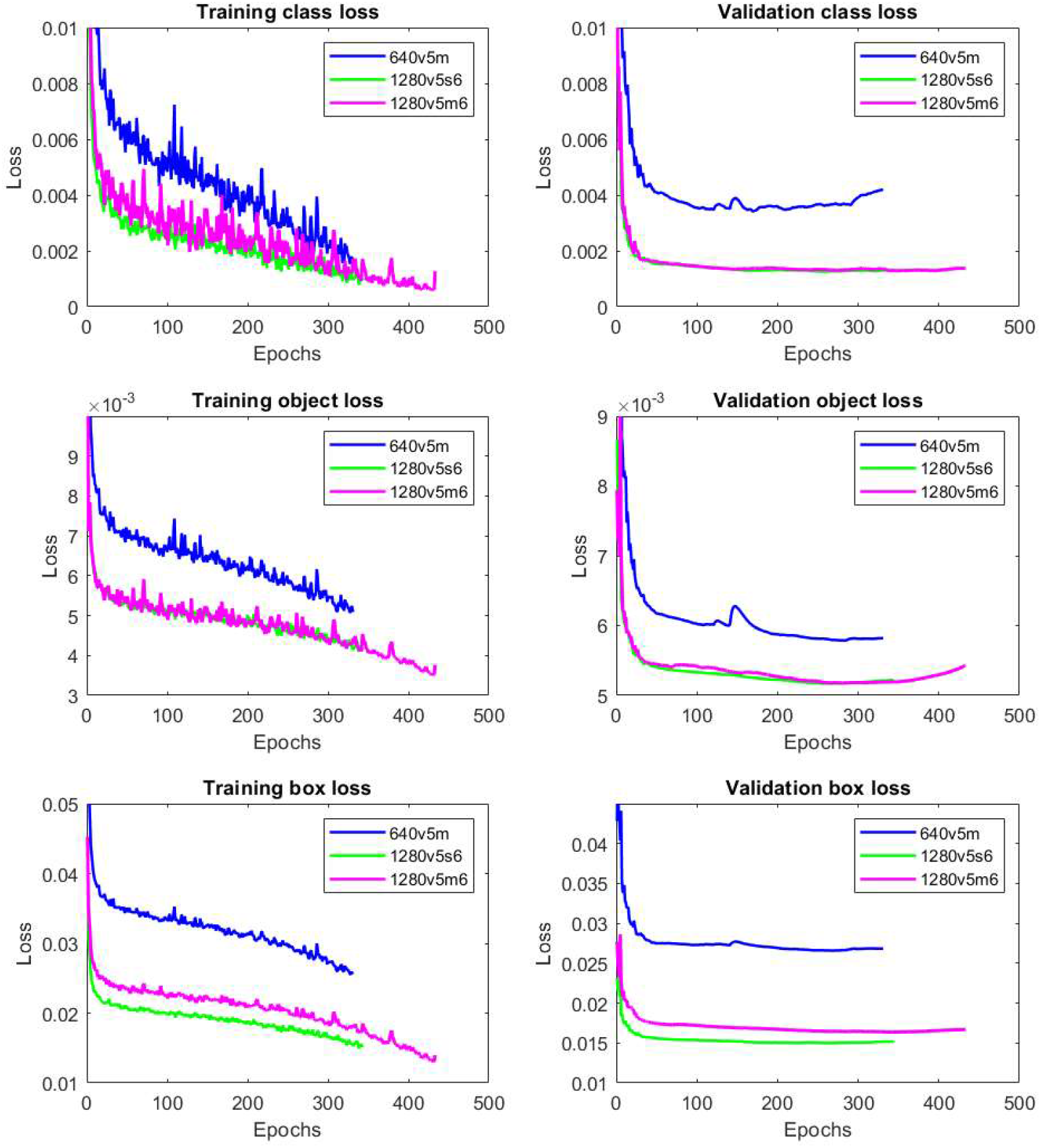
Training and validation with class, object and box loss for YOLOv5 models with nine classes. The loss is show as function of training epochs for three different YOLOv5 models: YOLOv5m (640v5m, blue), YOLOv5s6 (1280v5s6, green) and YOLOv5m6 (1280v5m6, red).

**Figure 5:**
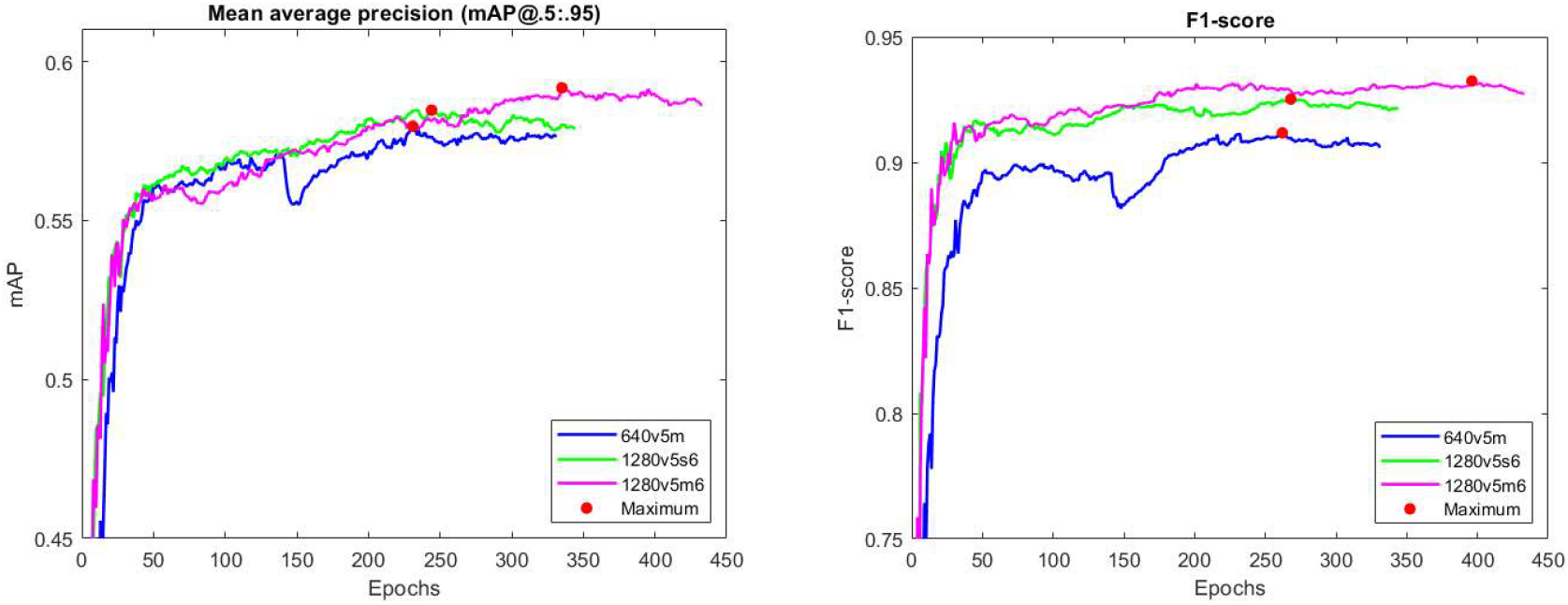
The mAP@.5:.95 and F1-scores are shown as a function of epochs for validating the trained models: YOLOv5m (640v5m, No. 5+6), YOLOv5s6 (1280v5s6, No. 7+8) and YOLv5m6 (1280v5m6, No. 9+10). The red marks indicate the best models based on the fitness function using the F1-score or the standard YOLOv5 fitness using mAP values. Training of 1280v5s6 (green curves) and 640v5m (blue curves) made an early stopping after 343 and 332 epochs since there was no increase of the fitness within the last 100 epochs.

The best YOLO models found are listed in Table *2* including one YOLOv3 model (No. 4). There was a remarkable improvement with the use of any of the configured YOLOv5 models compared to YOLOv3 model trained on the benchmark dataset. The large dataset, in particular, provided greatly improved performance of YOLOv5 models with an increased F1-score of 0.2 compared to training on the small dataset. This was not the case for YOLOv3. The increased image and model size from 640 (640v5m) to 1280 (1280v5s6) pixels provides only a slight improvement with an increased F1-score of 0.01 with nine classes. The difference between the 1280v5s6 and 1280v5m6 models was also minimal with an increased F1-score lesser than 0.01. The method of using the F1-score for the fitness function gave only a slightly better result seen on the F1-score. The absolute best YOLOv5 model was no. 10 with an F1-score of 0.932.

The best-trained YOLOv5 model is shown in Table *3* with metrics for each of the nine classes of species. The most common class in the images was the honeybee (*Apis mellifera*), which had a very high precision of 95%. The most difficult species to detect and classify were *Eristalis tenax* and *Episyrphus balteatus*; 5% of *E. balteatus* were detected as background false negatives. For *Eristalis tenax* the amount of training (286) and validation (28) data was limited, and this species is easily mistaken for, and widely considered to be a Batesian mimic of, *Apis mellifera* (19). This is reflected in the model confusion matrix in Figure *6*; 18% of *Eristalis tenax* was predicted as *Apis mellifera*. A high recall and precision was achieved for *Aglais urticae*, even with few training data. The confusion matrix indicates a good model in detection and classification of species with a diagonal of high scores ranging from 0.82 to 1.00. However, there are a number of false positive (FP) detections in the background image, especially for *Apis mellifera* (0.44) and hoverflies (0.36).

**Figure 6:**
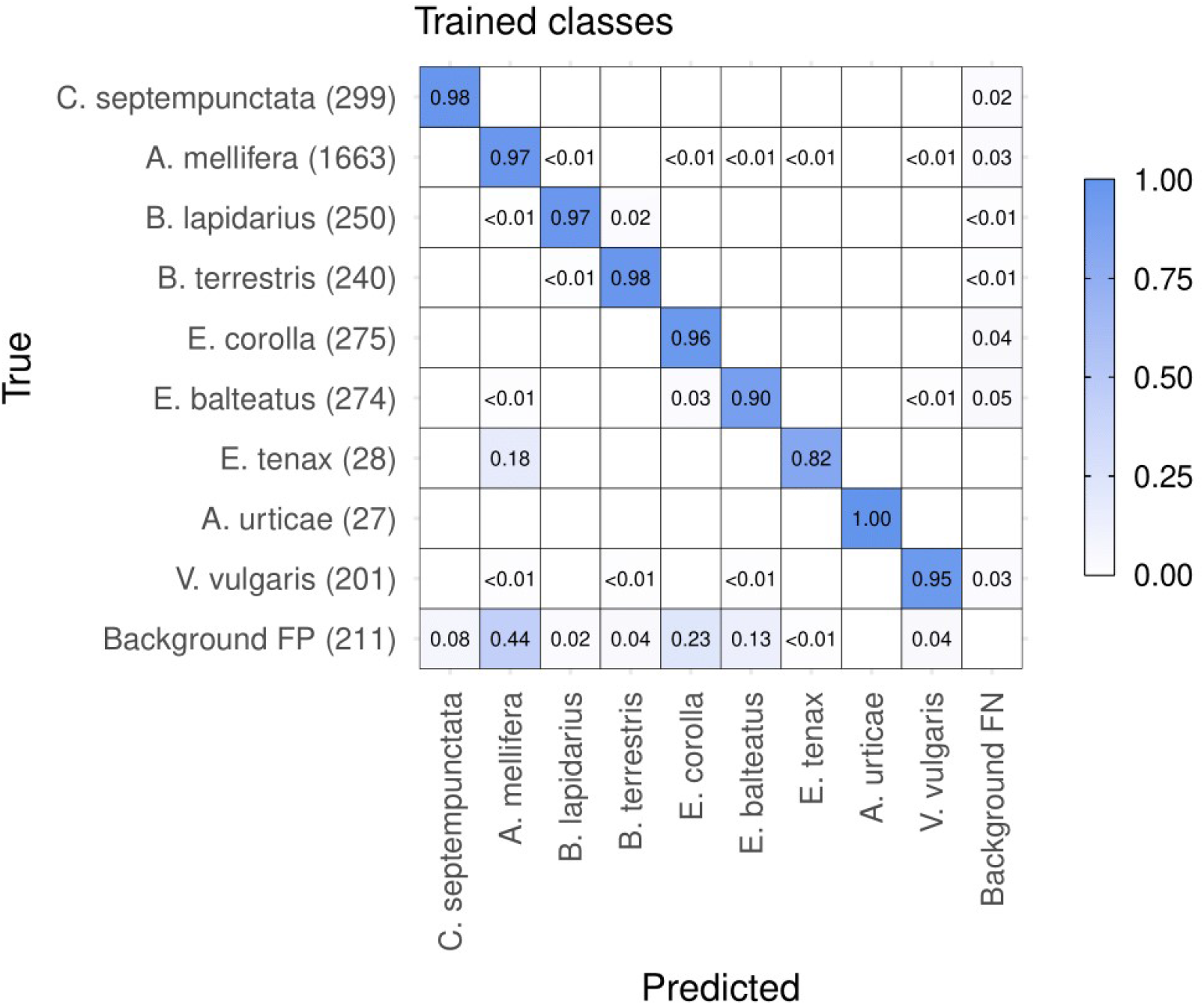
Normalized confusion matrix of predictions of the best YOLOv5 model (No. 10) for trained classes within the validation dataset. The y-axis (True) is the ground truth class of annotated insects. The number of observations of each class is shown in brackets. The x-axis (Predicted) is the class predicted by the trained model. The values in cells indicates the proportion of each true class that was attributed a given predicted class. For example, 82% of *Eristalis tenax* were predicted correctly, with 18% falsely classified as *Apis mellifera*. 44% of 211 false positive detections in the background image were predicted as *Apis mellifera*.

To evaluate whether the best model will detect unfamiliar insects and interpret them as closely related species, predictions were made for the additional test dataset using the trained YOLOv5 model (Table 2, No. 10). Of 4,212 unfamiliar insects, 3,509 (80%) were detected, of which most were classified as closely related taxa (Fig. 7). In particular, unfamiliar classes which were represented in the training dataset by multiple species in the same genus or family (e.g. Bombus spp., Syrphidae) were detected and classified in a sensible way (Fig. 7). Most *Bombus* spp. were detected and classified as either *Bombus lapidarius* or *Bombus terrestris* (i.e. Bombus spp., 91%, Fig. 7), while Syrphidae were quite consistently detected classified as *Eupeodes corolla*, *Episyrphus balteatus* or *Eristalis tenax* (80%). Indeed, the vast majority of non-*Bombus* Anthophila were appropriately detected and classified as *Apis mellifera* (92%). However, not all unfamiliar insects were so readily detected; Diptera were not detected in 48% of cases (Fig. 7). A total number of 684 or 7.8% false detections of plant parts (false positives) in images were mostly classified as honeybees and hoverflies, and are not shown in the matrix. A number of 997 or 11.3% missed detections (false negatives) gives an indication of how many insects will be missed in images recorded by the system.

**Table 2:**
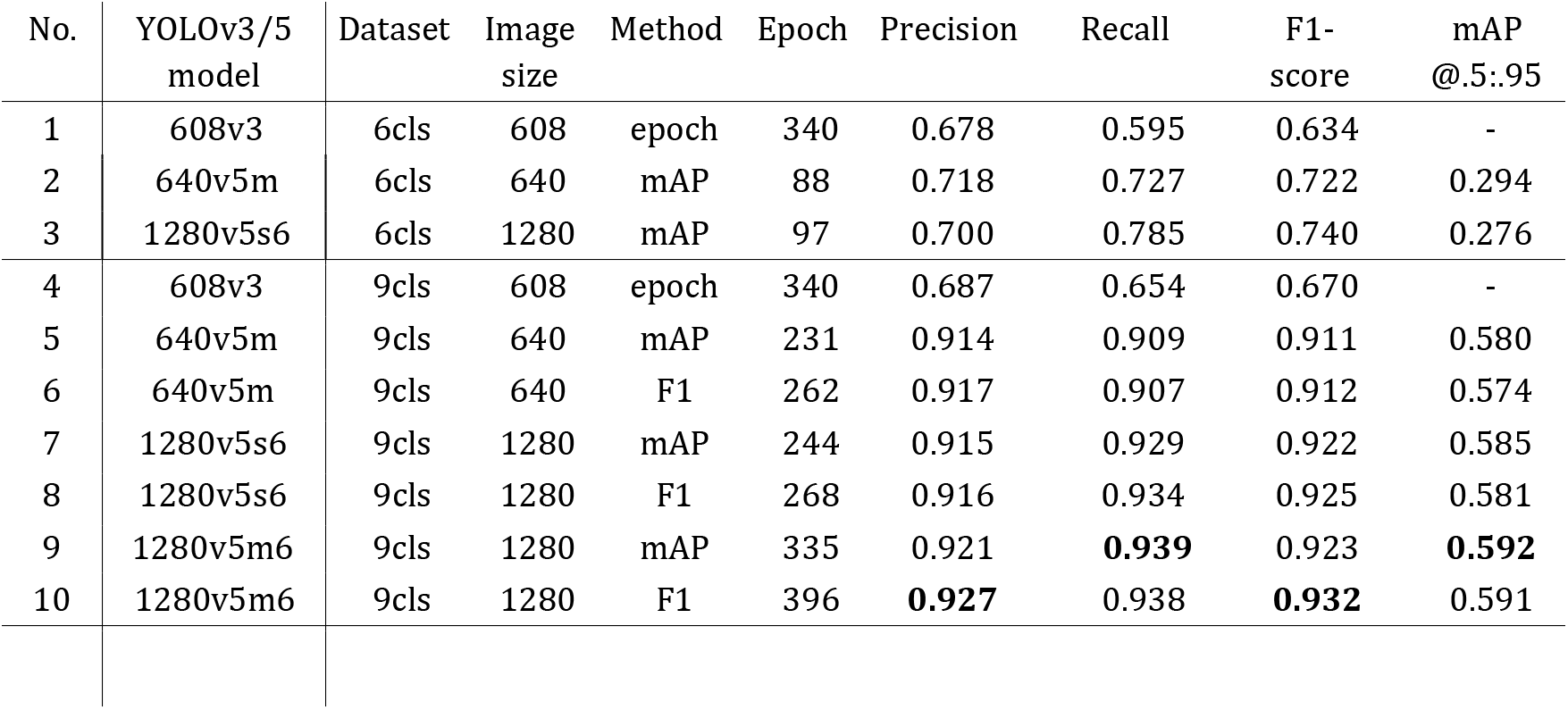
Precision, recall, and Mean Average Precision (mAP@.5:.95) on the validation dataset for different trained YOLO models. Models trained on the small (6cls) or large (9cls) dataset. The training method indicates which metric is used to evaluate when to stop the training process: mAP@.5:.95 (mAP), F1-score (F1) or stopping after a high number of epochs (epoch).

**Table 3:** Precision, recall, F1-score and mAP on the validation dataset of the best-trained YOLO model (No. 10) for each species in the dataset.

| No. | Species | Precision | Recall | F1-score | mAP<br>@.5 | mAP<br>@.5:.95 |
| --- | --- | --- | --- | --- | --- | --- |
| 1 | *Coccinella septempunctata* | 0.950 | 0.949 | 0.949 | 0.966 | 0.651 |
| 2 | *Apis mellifera* | 0.944 | 0.951 | 0.947 | 0.965 | 0.610 |
| 3 | *Bombus lapidarius* | 0.975 | 0.949 | 0.962 | 0.987 | 0.660 |
| 4 | *Bombus terrestris* | 0.959 | 0.975 | 0.967 | 0.982 | 0.648 |
| 5 | *Eupeodes corolla* | 0.882 | 0.949 | 0.914 | 0.938 | 0.504 |
| 6 | *Episyrphus balteatus* | 0.919 | 0.883 | 0.901 | 0.902 | 0.473 |
| 7 | *Eristalis tenax* | 0.799 | 0.854 | 0.826 | 0.853 | 0.605 |
| 8 | *Aglais urticae* | 0.963 | 1.000 | 0.981 | 0.966 | 0.675 |
| 9 | *Vespula vulgaris* | 0.949 | 0.935 | 0.942 | 0.938 | 0.502 |
|  | **Average** | 0.927 | 0.938 | 0.932 | 0.944 | 0.592 |

**Figure 7:**
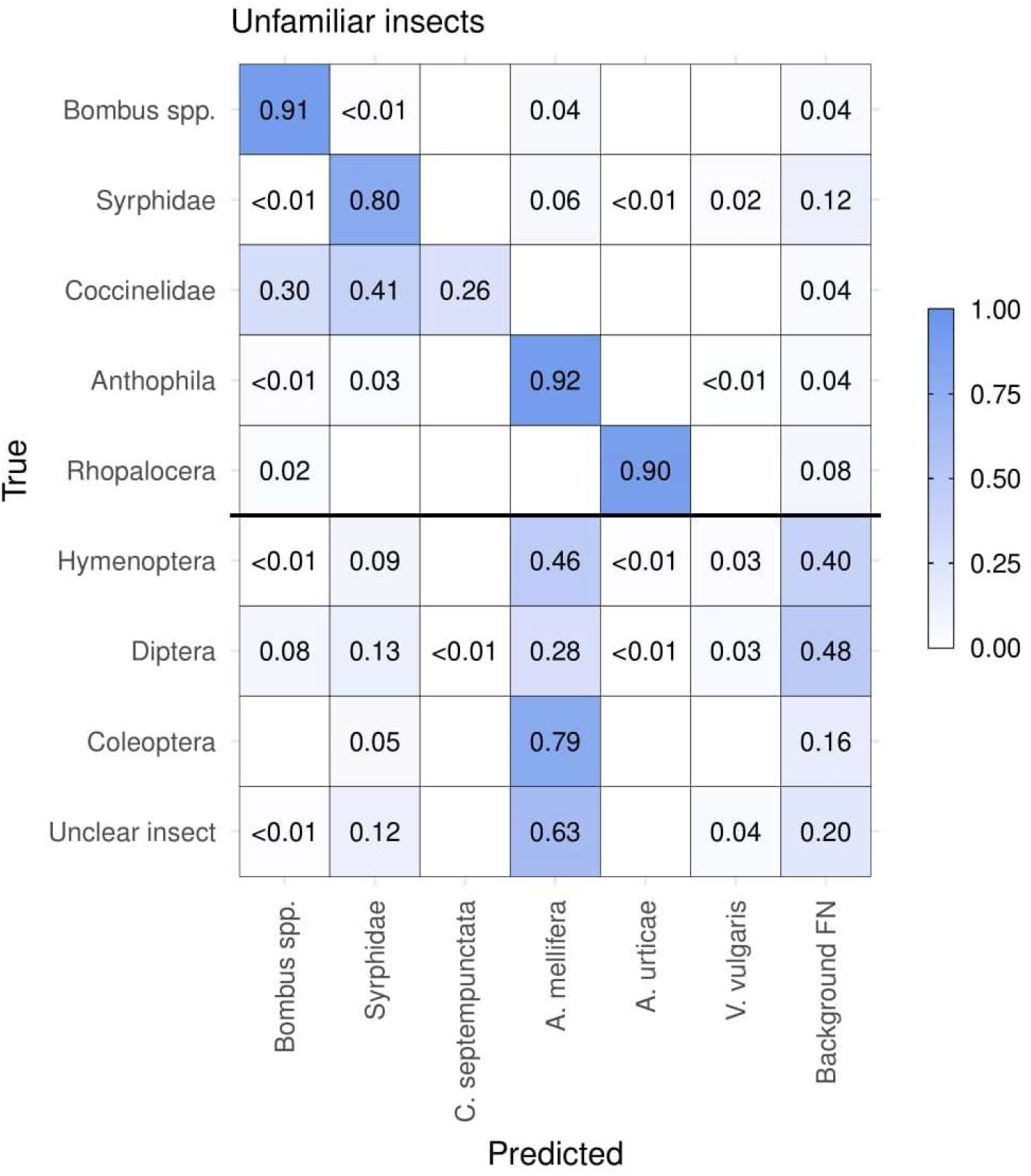
Normalised confusion matrix of predictions of unfamiliar insects present in the test dataset, based on the best YOLOv5 model. The y-axis (True) is the broad taxonomic class of unfamiliar insects. These unfamiliar classes only contain insects not assigned to any of the nine trained classes. The x-axis (Predicted) is the class predicted by the trained model, but aggregated trained classes were formed for Bombus spp. and Syrphidae for direct comparison with their unfamiliar equivalents. Detection and high-level classification accuracy was high for unfamiliar classes represented by at least one similar taxon during model training (above the solid black line).

All the collected images in the period from June to September from 10 cameras with a time-lapse of interval of 30 seconds between each image were finally analysed. Placed in a U-form, the cameras cover an area of approximately one square meter in total, as shown in Figure 1. Figure 8 shows the insect species detections per day found by the trained YOLOv5 model (1280v5m6) in the more than two million images recorded during the period of experiment. There is a clear seasonal dynamic in the occurrence of the various insects, while the occurrence of honeybees and hoverflies are more than 10 times higher than all other groups. It seems that hoverflies and ladybirds dominate the first half of the season (June-August), while the honeybees are most active in late August. The occurrence of *Eristalis tenax* was high in September, which deviates from other hoverflies (*Eupeodes corolla* and *Episyrphus balteatus*) most active in July. The variations in visit rates are be impacted by the weather and the longer-term seasonal effects such as different species of blooming sedum plants.

**Figure 8:**
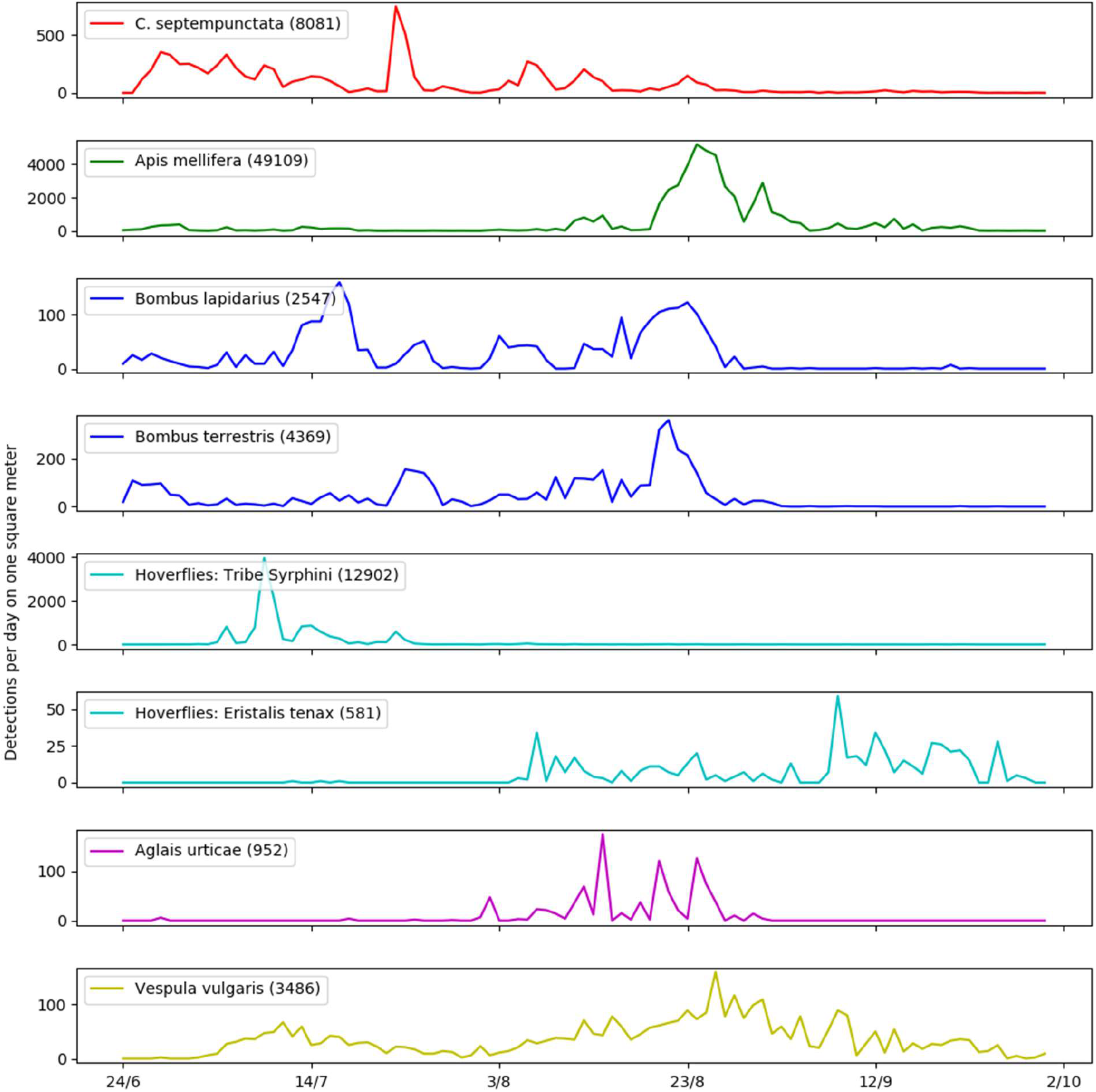
Seasonality and variation in the number of individual detections for each of the nine insect taxa counted in images collected from 10 cameras every day. The number in brackets shows the total count of the given species in the experimental period. Hoverflies holds the combined detections of *Eupeodes corolla* and *Episyrphus balteatus. Eristalis tenax* is separated in its own graph since it differs in abundance occurrence of the period.

## 3. Discussion

Insect detection in close range time-lapse photography is challenging as the objects of interest are small and varied and may resemble elements of the background (20,21). Earlier work has demonstrated that the amount of training data is critical to solving this challenge (22). With our study, we provide the tools and benchmark data to advance the implementation of robust automated monitoring of insects *in situ*. Our system is widely applicable as a means to capture images of plants and insects and to automatically generate monitoring data of insect species abundance (4,23–25). Our method could also contribute to insect pest monitoring with camera-equipped traps in agriculture and forestry without killing rare insect species (26,27). Remarkably, our automated insect monitoring camera system can collect images with sufficient resolution to efficiently and accurately analyse individual insects with YOLOv5. We have created and published a dataset with the most common insects in Europe for future training of deep learning models for automated insect detection and identification.

The best trained YOLOv5 model (1280v5m6 validation with highest F1-score) measured an average precision of 92.7% in classifying the species correctly with an average recall of 93.8 %. We know of no study with such high classification performance for insects recorded with time-lapse cameras, where image quality of the insects typically is lower than in portals such as iNaturalist. Comparable classification rates (91.5%) have been achieved in a similar study of insect classification and detection in field crops using machine learning techniques (28). However, this study did not tackle the challenge of detecting insects in larger images, and the image data sets (29) were not derived from time-lapse images. As image-based monitoring methods evolve, it will be important to evaluate how performance of classification models trained on different image data compare.

Image classification performance may vary among taxa. Our classification precision was poorest for hoverflies due to many false positives where parts of the background were detected as insects. Many of the false positives were structures in *Sedum* flowers, which appeared similar to the small hoverflies. *Episyrphus balteatus* is small compared to the other commonly detected insects in our data and could easily be missed. We also found that insects were misclassified. In particular, drone flies were sometimes misclassified as honeybees The appearance of the butterfly *Aglais urticae* is very distinct and different from the other insect classes in our data and this class had a high recall and precision even with little training data. Still, a high performance of the CNN supports our approach of incorporating computer vision and deep learning into the image recording process for insect monitoring. The improved and trained YOLOv5 model could easily replace the YOLOv3 model used in the real-time insect tracking and monitoring system presented by Bjerge, Mann, & Høye, 2021. We have measured that the YOLOv5s6 model takes less than 0.5 sec. to execute on the Jetson Nano computer, which means the frame rate increases from 0.33 fps (YOLOv3) to 2 fps. The newly released YOLOv7 (30) would be an alternative YOLO model to use, however initial results with YOLOv7-E6 does not improve performance on our benchmark dataset compared to YOLOv5 models presented in this paper.

Importantly, our YOLOv5 model, trained on nine insect species, is able to detect insects belonging to species that were not seen during training. Specifically, when presented with images containing unfamiliar or unclear insects, the model detected 80% of individuals. Thus, we raise an exciting prospect: The use of rich training data on a few common species to facilitate detection of rare or poorly documented insects. However, unfamiliar insects were most readily detected if many similar species featured in the training data (e.g. *Bombus* spp. and Syrphidae); thus, the model’s ability to detect novel taxa might improve if more species were featured in training. Fortunately, our model also represents a tool to efficiently generate training data for rarer insect species; misattributions of unfamiliar insects to familiar species can be manually screened. This would enable generation of ever-larger image reference libraries, spanning ever-more uncommon taxa. When algorithms and computer power develop sufficiently, we propose that future models employ a hierarchical classification process (31,32). This would incorporate the prospect of novel and nested taxa from the outset and generate predictions at the most appropriate taxonomic level.

Our camera system is composed of off-the-shelf hardware and supports sufficient storage space to work off-line in monitoring insects over the entire growing season. The system is a low-cost solution integrated with software based exclusively on open source code. The solution operated with only one interruption over extended periods, even during rain, strong winds, and at high temperatures. During our experiment, we ran the deep learning algorithm after image recording was completed, but it could be moved to on-site processing on a Raspberry Pi 4B, Jetson Nano or similar. By running the entire algorithm on the camera system, it would also be possible to present summary data in real-time to the end-user of a specific unit or to combine summary data from multiple units at a regional or even global scale to a wider community. Once the system is launched, it requires little to no maintenance, and communication via the internet provides an opportunity to monitor the status of each system. The total cost of the hardware with Logitech C922 web cameras, Raspberry Pi 3B and an SDD hard drive is approximately $350 with grid power. Solar panel, battery and charger add an additional cost of $425 for locations without power. The cheaper plant-insect interaction camera trap (PICT) is based on a Raspberry Pi Zero with only one camera and storing data on a micro SD card (33). The PICT system could be modified with Logitech C922 cameras to lower the price. As such, we argue that our solution is tested, stable, reasonably priced and thus ready for large-scale deployment for insect monitoring across larger regions.

The proposed trained YOLOv5 solves an important challenge for insect monitoring by applying CNN-enabled object detection after the recording process. However, the models are trained on a time-lapse images from 10 cameras with very similar backgrounds, thus model performance will almost certainly be lower on images with other plants and flowers in different contexts. We present an important benchmark dataset to train models for insect detection and classification tasks. However, to achieve better generalisation performance, this dataset must be combined with other, new datasets from a wide variety of environmental contexts. In future, generalisation performance might be rigorously tested by holding out specific cameras or contexts during model training. Furthermore, we have presented a single-pass detection and classification process using YOLO. Alternatively, a two-pass approach could be proposed starting with a general insect detector followed by a higher performance classifier to identify insect species. This two-phase approach might generalize better to other contexts if the insect detector was trained with a higher variation of background images. It might also cope better with unbalance in class frequencies, especially during detection.

If a model is applied after data collection, with the same camera system as presented in this paper, the model will likely be able to detect the nine different insect species. Some of the unfamiliar species will be detected, but they would be classified as one of the nine species in the dataset. A strategy to handle new species would be to extend the dataset with more classes of species and retrain the YOLOv5 model. In a novel environment background images needs to be added to the dataset and the model retrained for it to learn and distinguish insects from plants in the new background. There are only minor difference in the precision and recall for the best trained YOLOv5 models, so for on-board processing either the YOLOv5m, YOLOv5s6 or YOLOv5m6 models should be considered since speed is important for real-time detection. The proposed computer vision system and deep learning with YOLOv5 is promising as an automated system for insect observations and complements existing systems for monitoring nocturnal insects (34).

## 4. Materials and methods

### 4.1. Collection of images

Images were collected using five recording units each consisting of a Raspberry Pi 3B computer connected to two Logitech C922 HD Pro USB web cameras (35) with a resolution of 1920×1080 pixels. Images from the two cameras were stored in JPG format on an external 4TB USB hard disk (Figure *1*). The Raspberry Pi 3B with two cameras and a hard drive consumes only 4W during full operation powered by 5V. We supplied power to the system through the grid connection (230-240V AC) connected to a 5V/2.5A transformer, but due to the low power consumption, each unit could be powered by a 12V 55Ah lead battery, DC-DC converter and a 60Wp solar panel for deployment at locations without power. A time-lapse program (36) installed on the Raspberry Pi was used to continuously capture time-lapse images with a framerate of 30 seconds between images. This program could also be configured to trigger whenever movement is detected in the camera field of view. However, it is not guaranteed that insect movements are sufficient, while movement of plants would create many images without insects. The camera used automatic exposure to handle variation in light conditions related to direct sun, clouds, and shadows. Focus was manually adjusted for optimal image sharpness. With continuous time-lapse recording of 30 seconds, we recorded a maximum of 2040 images per camera per day between 5 AM and 10 PM in the timespan when there was sufficient light to capture images

During the period June to September 2019, the camera systems were in operation on a green roof in Aarhus, Denmark (GPS Latitude 56.153471, Longitude 10.212053) with a mix of 13 species of *Sedum* plants (18). Four cameras were adjusted to a working distance of 24 cm covering an area of 35 × 22 cm and six cameras with a working distance of 32 cm covering an area of 46 × 27 cm. An insect (0.5 – 2 cm) occupies between 45×45 to 120×120 pixels and the captured images were stored in JPG format with a resolution of 1920×1080 pixels.

### 4.2. Benchmark insect image datasets

To efficiently generate a large benchmark dataset for detection and classification of insects, we used a three-stage approach. In the first stage, insects were annotated within a subset of images to train a preliminary deep learning model. In the second stage, predictions from the preliminary model provided many more candidate insect detections, which were verified and used to generate improved deep-learning models. In the third stage, to evaluate model responses to unfamiliar classes, uncommon or unclear insects were annotated to a coarse taxonomic level.

In the first stage images were selected opportunistically from the ten cameras, capturing variation in insects, flowering *Sedum* plants from different cameras, and the full growing season. In the images with insects, we made bounding box annotations, using a labelling image tool (37), of what appeared to be the six most abundant insect taxa (*Apis mellifera*, *Bombus terrestris*, *Bombus lapidarius*, *Episyrphus balteatus*, *Eupeodes corolla*, and *Coccinellidae septempunctata*). More than 1,000 background images without any insects were also selected within fixed time and date intervals from all 10 cameras. The resulting images across all cameras were randomly assigned to either training or validation datasets to train a preliminary YOLOv3 model as described in (18). This preliminary model located a large number (127,171 detections or candidate insects) of additional insects in the more than two million recorded images.

In the second stage, the preliminary model was used to make predictions across all images recorded by the 10 different cameras during the whole recording period. These predictions generated a large number of images with candidate insects, which were verified for the final benchmark dataset as follows. First, candidate insects were manually screened by users of our online citizen science project PollinatorWatch on the Zooniverse platform (https://www.zooniverse.org/). Subsequently, candidate insects were quality checked by the authors (T. Høye) and bounding boxes were adjusted especially for butterflies. Finally, the working taxonomy was updated to account for new classes with high representation in the data, for a total of nine classes (see Table 1). During quality checks, non-detected insects were found in the same images as detected and verified insects; these were annotated and included in the benchmark dataset. Finally, images with annotations were separated in a stratified-random way into three groups (training, validation, and test data). Images were separated to provide even representation over time of day and season across all 10 cameras. For the benchmark dataset a split of approximately 85/15 was chosen to prioritize more data for training, but still having sufficient data samples to validate the models. The validation dataset contained at least 200 annotations of each species except for *Aglais* urticae and *Eristalis tenax*, for which 10% of the annotations were used for validation.

Finally, uncommon or unclear insects were annotated. Throughout verification, many insects had unclear identity or fell outside of the nine training classes. These organisms were not considered useful for training, but they represent an important part of the reality of automated *in situ* insect monitoring. To understand how a trained model might behave when presented with novel or unclear insects, we produced an extended test dataset including nine additional classes of uncommon or unclear insects, as well as other animals (e.g. slugs; Table 1). The additional insect classes comprised unknown bumblebee species (*Bombus* spp.), hoverflies (Syrphidae), ladybirds (Coccinelidae), non-*Bombus* bees (Anthophila), butterflies (Rhopalocera), non-bee Hymenoptera (mostly sawflies: Symphyta), non-Syrphid flies (Diptera), non-ladybird beetles (Coleoptera) and unclear or blurry insects (usually in motion or out of focus). We refer to these as unfamiliar classes, as they were completely excluded from the foreground and background of both the training and validation sets.

### 4.3. Insect detection and classification with YOLO

To find insects of interest, different versions of the deep learning algorithm YOLO (17) were trained and evaluated. YOLO is a one-stage object detector, which predicts bounding boxes around objects (insects) within the image, their class labels (species) and confidence scores. One-stage object detectors are faster than two-stage detectors such as Faster R-CNN (38) with the cost of lower accuracy. However, fast execution is important when millions of images need to be processed, or the object detector is deployed on an embedded computer *in-situ* e.g. (14). The collected images were processed by YOLO on a workstation computer in our experiment.

In this paper, YOLOv3 and YOLOv5 architectures were evaluated. YOLOv3 was used to produce the preliminary model, as it was accessible when we started the study in 2019. Later YOLOv4 and recently YOLOv5 were published, with many new features, and we evaluate this evolution of models in detecting and classifying insect species. YOLOv3 was used with the CNN backbone darknet53 with a total of 53 layers, which has achieved a high mAP on the benchmark COCO dataset of 57.9 % (39). This accuracy was comparable to other state-of-the-art deep learning networks like RetinaNet (8) and ResNet50-based R-CNN (40). Darknet53 uses three residual skip connections that make detections at three different scales. YOLOv4 (41) uses the modified backbone CSPDarknet53 and introduces new features such as “Bag of specials” and “Bag of freebies” where the data augmentation for training is improved without affecting the inference cost. “Bag of freebies” covers a number of new augmentations such as CutMix and Mosaic. Mosaic mixes four different training images as opposed to CutMix using two images to enhance the data augmentation. In order to tackle class imbalance in our dataset, a focal loss function is used (42) in YOLOv4. The focal loss function consists of a dynamical scale cross-entropy loss where the scaling factor is adjusted based on the confidence. This means that the scaling will decrease towards zero as the confidence increases. As such, the easily recognisable insects have less impact on the training. The activation function in YOLOv4 is changed to the novel self-regularizing non-monotonic activation function (Mish) (43). YOLOv5 (44) is developed in parallel with YOLOv4 and contains the same features implemented natively in Python and PyTorch. YOLOv5 is easier to train as it introduces auto-anchor. Before auto-anchor, the anchor boxes would need to be trained as a step before training. YOLOv5 provides capability of models with different neural network architectures and image sizes. The smaller models allow faster execution. To improve performance and speed up training, the YOLOv5 is pre-trained on the COCO (39) dataset. We used the pre-trained models and compared the capability of YOLOv5m, YOLOv5s6 and YOLOv5m6 to detect and classify insect species. The models were chosen because their size allows them to be run with the required memory, and achieving a high accuracy on most reasonably priced GPUs. An official paper for YOLOv5 has not been released, but YOLOv5 has been successfully used to study the abundance of three pest insect species on sticky traps (45).

In the training of YOLOv5, three different loss functions were calculated. The first concerns errors between the predicted boundary of the bounding box and ground truth (box loss). The second is confidence in whether an object is detected in the box (object loss). The third, classification loss, concerns if an object is correctly classified (class loss) and is the sum of squared error of the class conditional probabilities for each class.

To evaluate the potential trade-off between model performance and processing speed with variable image sizes we explored different combinations of image size (640×384 and 1280×768 pixels), small (YOLOv5s6 with 12.6M parameters) and medium (YOLOv5m with 21.2M parameters, YOLOv5m6 with 35.7M parameters) model size, and two different fitness functions to find the optimal YOLOv5 model to classify the insect species in our data. The accuracy of the YOLOv5 models evaluated in this paper are ranging from 45 to 51 in mean Average Precision (mAP) on the COCO dataset (44). The default fitness function in YOLOv5 defines a weighted combination of metrics, where mAP@.5 contributes 10% of the weight and mAP@.5:.95 contributes with the remaining 90%. mAP@.5 is the mAP calculated as the mean area under the precision-recall curve for all classes with an IoU of 0.5. mAP@.5:.95 is defined by the COCO challenge (39) as the average mAP across all the IoU threshold between 0.5 and 0.95 with a step of 0.05. Here, we have compared this default method with using the maximum F1-score as the fitness function. The F1-score is calculated based on the recall and precision as:

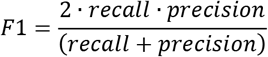

The F1-score prioritizes the importance of detecting and classifying the insect species correctly over fitting the correct bounding box size. Since insects are very small in the images there is a high uncertainty of the annotated bounding box.

To demonstrate the value of automated cameras and to understand insect phenology, we present the number of occurrences per day of different insect species throughout the period of the experiment. To better understand the role of YOLO model generation and image size for model performance, we also trained a YOLOv3 model using default settings trained with an image resolution of 832×468 pixels and validated on images with 608×342 pixels. This approach gave better results than training on images with 608×342 pixels (18). The number of training iterations for the YOLOv3 model was chosen based on when the average training loss was no longer decreasing. The selected YOLO3 model used 30,000 iterations (equal to 340 epochs with a batch size of 64), which gave a higher precision and recall on the validation dataset than using less than 28,000 iterations. A higher number of training iterations would increase the risk of overfitting the model.

## Acknowledgments

The work was supported by the European Union’s Horizon 2020 Research and Innovation programme, under Grant Agreement no. 773554 (EcoStack) and no. 101060639 (MAMBO).

## Author contributions

K.B., J.A., H.M.R.M. and T.T.H. conceived the ideas and methodology; J.A. and T.T.H conducted the annotation and validation of datasets; K.B., J.A., and T.T.H. performed the analyses of results and production of figures; K.B. trained and validated the YOLO models and led the writing of the manuscript; M.D., C.E.F. and H.M.R.M. provided valuable recommendations for improving results. All authors contributed critically to the drafts and gave final approval for publication.

## Data availability statement

The benchmark and test dataset can be downloaded from: https://doi.org/10.5281/zenodo.7395751 or https://vision.eng.au.dk/ecostack

